# Sunflower yield modeling with XAI: Historical weather impacts and forecasting

**DOI:** 10.1101/2025.02.27.640573

**Authors:** Sambadi Majumder, Chase M. Mason

## Abstract

This study applies explainable artificial intelligence (XAI) to analyze the impact of inter-year variation in weather conditions on oilseed sunflower yields across the United States. By integrating historical yield data from 1976 to 2022 with meteorological data, we identified key weather predictors influencing sunflower yields at national and state levels along with critical yield-sensitive threshold temperature and precipitation values that predict reduced yield. Using machine learning algorithms, we developed predictive models to forecast future yields under different Shared Socioeconomic Pathways (SSPs) from 2021 to 2080. Our findings reveal significant yield declines due to increased temperatures and altered precipitation patterns, but with regional variability in the magnitude of these impacts. The most critical climate variables identified include maximum temperatures and total precipitation during summer months. Our XAI approach enhances model transparency, offering valuable insights for farmers and policymakers to develop adaptive strategies for sunflower cultivation under climate change. Future research incorporating additional factors like soil characteristics and agricultural practices can further refine yield predictions.

## 1. INTRODUCTION

Adverse impacts to global food production are predicted to occur under global climate change (Iizumi et al., 2015; Siebert et al., 2017), with the possibility of large-scale crop losses due to extreme weather at a range of scales (Mehrabi et al., 2019; Tigchelaar et al., 2018). Accurate forecasting of the expected impacts of climate change on crop yield at a fine spatial level would enable farmers and policy makers to evaluate current farming practices and facilitate future planning in support of food security (Mann et al., 2019). Yield is an important metric of agricultural production and forecasting changes in yield is imperative to aid management of food availability under climate change. Yield is influenced by non-additive and nonlinear interactions of crop genotype, environmental conditions, and management practices, making the accurate prediction of yield changes a complex problem (Horie et al., 1992; Khaki et al., 2020). Machine learning (ML) approaches have proven to be extremely successful in predicting crop yields, in large part due to the ability of such models to handle complex nonlinearity in the data (Khaki et al., 2020; van Klompenburg et al., 2020), and the ability to incorporate different data types in the modeling process, for example remote sensing (Ahamed et al., 2015; Fernandes et al., 2017; Shahhosseini and Archontoulis, 2020; Qiao et al., 2021), image data (Črtomir et al., 2012; Cheng et al., 2017) and spatial climate data in raster formats (Mola-Yudego et al., 2016). Researchers using ML can utilize information from various data sources and formats and thus include information on crop progress, environmental conditions, and their relationship with yield, resulting in reliable and accurate estimations (Han et al., 2020; Shahhosseini and Archontoulis, 2020; Zhang et al., 2020). This has led to widespread applications of ML in “precision agriculture” both for crop yield forecasting and as a data mining tool (Ruß et al., 2008; Everingham et al., 2009; Ahamed et al., 2015; Gandhi et al., 2016). Examples include predicting yield in major commodities like corn (Kang et al., 2020; Shahhosseini and Archontoulis, 2020), soybean (Habibi et al., 2021;Yoosefzadeh-Najafabadi et al., 2021), coffee (Kouadio et al., 2018), wheat (Pantazi et al., 2016; Cai et al., 2019; Li et al., 2022; Liu et al., 2022), sugarcane (Everingham et al., 2009), and rice (Gandhi et al., 2016; Su et al., 2017; Cao et al., 2021; Son et al., 2022). The incorporation of interpretable methods in ML modeling makes it possible to disentangle the complex relationships that exist in spatiotemporal multidimensional agricultural datasets (Mateo-Sanchis et al., 2021; Ryo, 2022). The emerging field of explainable AI (XAI), specifically interpretable ML, offers tools to enhance the interpretability of complex algorithms without sacrificing predictability (Doshi-Velez et al., 2017; Adadi et al., 2018; Molnar, 2019; Murdoch et al., 2019; Rudin, 2019). XAI has gained attention and potential in various natural science fields, including biodiversity research (Ryo, 2022), geoscience (Mamalakis et al., 2022), and hydrological/climatic science (Başağaoğlu et al., 2022). In the agricultural domain, XAI techniques have been applied to crop yield estimation (Sihi et al., 2022; Wolanin et al., 2020), crop type and trait classification using satellite data (Newman et al., 2021; Orynbaikyzy et al., 2020), soil texture classification (Zhou et al., 2022), leaf disease classification (Wei et al., 2022), water assessment (Garrido et al., 2022), smart agriculture systems (Sabrina et al., 2022), biomethane production (Clercq et al., 2020) and agricultural land use prediction (Viana et al., 2021). XAI can achieve a level of explainability alongside high accuracy that is essential for understanding predictions, evaluating their significance, and enabling informed decision-making (Mateo-Sanchis et al., 2021; Ryo, 2022). In agriculture, interpretable models can help identify the key variables influencing crop yield, assess the impact of their changes during the growing season, and detect anomalies and extremes in specific areas. Interpretability also aids in model evaluation, as it is imperative that those developing a model for “real world” application ensure that the model has learned meaningful patterns from the agricultural data and that the explanations behind each prediction align with existing domain knowledge (Mateo-Sanchis 2021; Ryo, 2022). Our study focuses on crop sunflower (*Helianthus annuus*), one of the most important oilseed crops globally (Mandel 2011; Seiler et al., 2017; Zymaroieva et al., 2021), and one of the few major commodity crops native to the United States (Heiser et al., 1969; Rieseberg et al., 1990; Crites 1993; Harter et al., 2004). Despite being a moderately drought tolerant crop generally adaptable to a wide variety of agroecological conditions (Miladinović et al., 2019), sunflower production is negatively impacted by climate change (Debaeke et al., 2017; Hussain et al., 2018; Tariq et al. 2018). Variation in the availability of water, both as precipitation and soil moisture, lead to inter-year differences in sunflower yield (Aboudrare et al., 2006; García-López et al., 2014). Heat and drought stress during certain growth stages like germination, anthesis, and seed filling has been shown to lead to approximately 50% decline of overall yield and oil quality in sunflower (Hussain et al., 2018; García-López et al., 2014). Increasing temperatures can also disrupt crop phenology, with mismatch between photoperiod and thermal regimes shifting the timing of flowering and seed maturation to less optimal periods and reducing yield (Ahas and Aasa, 2006; Cleland, 2007; Forrest and Miller-Rushing, 2010; Li et al., 2014; Hatfield et al., 2015). As extreme weather events are expected to become more common in the future (Iizumi et al., 2015; Siebert et al., 2017), researchers have used different forecasting strategies to accurately predict regional sunflower and other crop yields under climate change – though often with limited data availability (Wu et al., 2007; García-López et al., 2014; Iizumi et al., 2017; Yeşilköy and Şaylan, 2021). Scientists have also attempted to incorporate socioeconomic shifts when assessing the impact of climate change on agriculture (Nelson et al., 2014a, 2014b, von Lampe et al., 2014; Wiebe et al., 2014), given the critical impacts that population growth, rising global incomes, and increasing food consumption have had on the demand for land and its resources over the past century – triggering significant land use change and contributing to climate change (Goldewijk et al., 2011; Doelman et al., 2018). Researchers have examined the global and regional impacts of climate change on agriculture by combining multiple models to explore how different socioeconomic emissions pathways affect these impacts (Rosenzweig and Parry, 1994; Parry et al., 2004; Nelson et al., 2010; von Lampe et al 2014; Wiebe et al., 2014).

In this study, we identify meteorological variables that influence inter-year variation in sunflower agricultural yield per acre at both national and regional scales using interpretable ML approaches that identify critical yield-sensitive thresholds of temperature and precipitation during the growing season. In addition, we use these inferences to forecast average sunflower yield per acre at a county level across the United States for three separate future time periods (2021- 2040, 2041- 2060, and 2061-2080), assuming recent values of county-specific area planted and harvested and taking into consideration different socioeconomic carbon emissions scenarios (Riahi et al., 2017). We first identify specific climate variables that have historically influenced overall yield across seven states in the contiguous United States, and then describe how states vary in the relative importance of different climate variables. This modeling further describes the response of yields to inter-year variation in weather conditions and identifies potentially important agronomic thresholds for sunflower crop stress during the growing season.

## 2. METHODS

### 2.1. Software and programming language

Data was compiled, cleaned, and processed using a combination of the R programming language (R Core Team 2023) and ArcGIS Pro software (version 2.9), with all visualizations and modeling performed using packages within the R programming environment. All code to replicate the analyses in this study can be accessed through the project GitHub repository [**REDACTED FOR BLINDED REVIEW**]. Given the complexity of the machine learning modeling performed, an expanded methods section is provided with additional analytical details and accessible explanations of modeling steps (Supplementary Methods).

### 2.2. Study area, data sources, and data preparation

County-level annual “oil type” sunflower seed yield data (measured in pounds per acre) as well as accompanying area planted and area harvested (both in acres) were acquired programmatically from the United States Department of Agriculture (USDA) National Agricultural Statistics Service (USDA National Agricultural Statistics Service, 2017) using the R package *tidyUSDA* (Lindblad 2022) through an API key. This yield data spanned the temporal range between 1976 and 2022, and included 189 counties across North Dakota, South Dakota, Kansas, Nebraska, Colorado, Minnesota, and Texas. Historical monthly weather data was acquired from the National Oceanic and Atmospheric Administration (NOAA) National Centers for Environmental Information database (NCEI) as multi-layered netCDF raster files containing 5km gridded monthly maximum temperature, minimum temperature, and total precipitation records for the period 1976 to 2022 (Vose et al., 2023).

To obtain future climate projections, multi-layered raster files were obtained from the WorldClim 2.1 repository released in January 2020 (Fick et al., 2017) using the *geodata* R package (Hijmans et al., 2022). Downsampled data from the CMIP6 climate model ACCESS-ESM1-5 (Ziehn et al., 2020), at 2.5 arc-minute resolution (approximately 4.5 km grid cells) was selected to provide similar spatial resolutions for historical and future climate data. The future projections obtained reflected four Shared Socioeconomic Pathways (SSPs; Riahi et al., 2017): SSP 1-2.6 (sustainability), SSP 2-4.5 (middle of the road), SSP 3-7.0 (regional rivalry), and SSP 5-8.5 (fossil-fueled development) across three time periods (2021-2040, 2041-2060, and 2061-2080). Both historical and future weather raster files were first reprojected to the same coordinate system - USA Contiguous Albers Equal Area Conic Projection (Snyder, 1987) within ArcGIS Pro. This was followed by extracting data from each raster layer at the county centroids (National Weather Service, 1995) and storing the extracted data in a dataframe format. Further data preparation was performed within the R programming environment using the packages within the *tidyverse* suite (Wickham et al., 2019). Missing data was removed, and the dataset was divided into a training and a testing dataset using a temporal cut-off point of 2005 (Figure 1). Individual state-level training and testing datasets were also created using the subset of counties from the combined dataset within each of the seven focal states, with state-level training datasets contained 70-80% of the total data.

**Figure 1.**
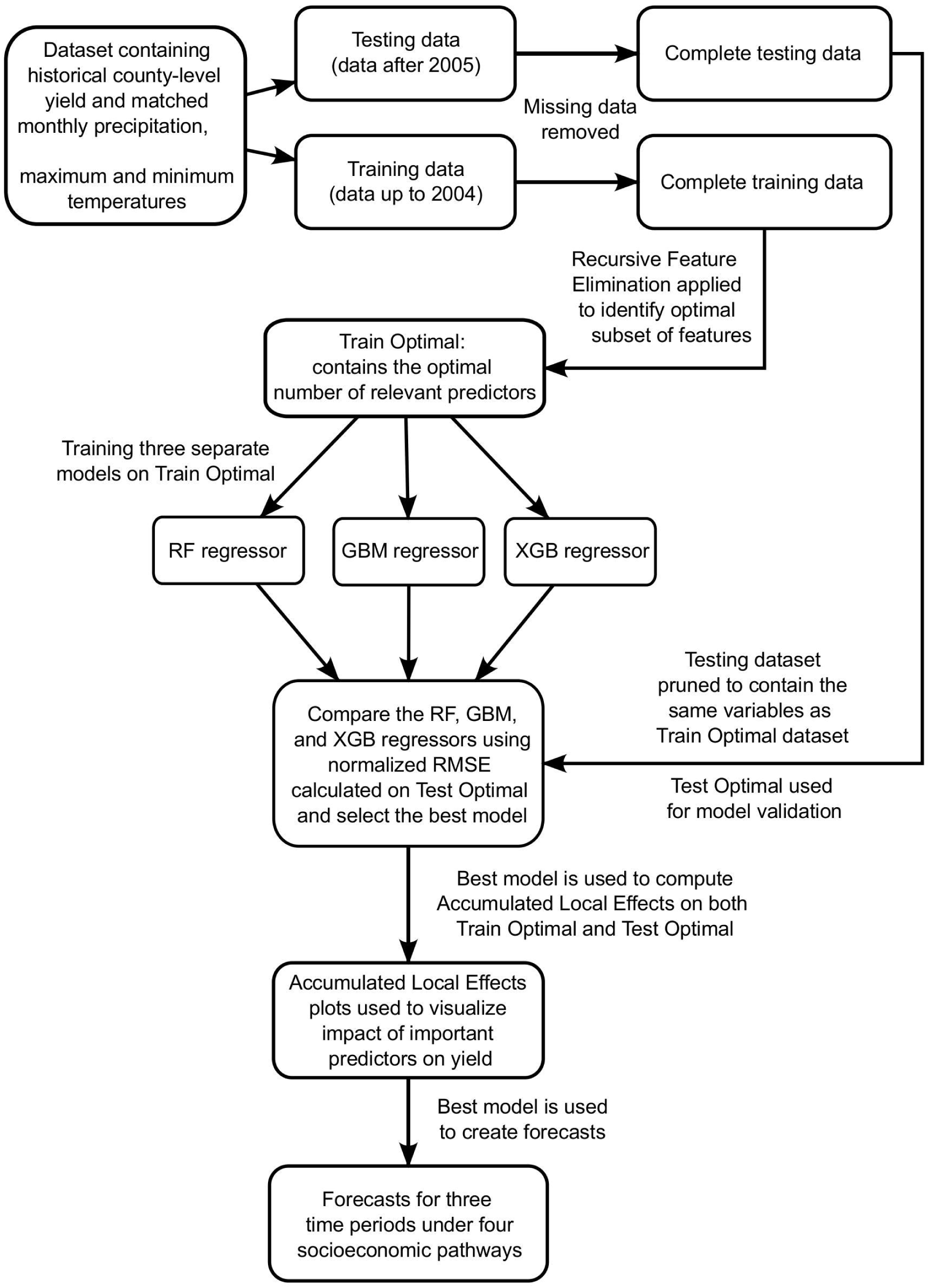
The complete workflow of the entire analysis procedure. This workflow was applied to both national dataset and state-level datasets which contained historical gridded (approximately 5km^2^) monthly precipitation, maximum and minimum temperature (predictors). The datasets also contained the total area planted and harvested (in acres) as well as historical yield data (measured in lbs/acre). The national dataset contained data from all 177 counties used in this study, while the state-level datasets only contained data from counties in that given state. Both types of datasets (national and state level) were temporally divided into training and testing datasets. The temporal cut-off point used in each case was the year 2005, where data up to 2004 was used as the training dataset and data from 2005 onward was used as the testing dataset. A recursive feature elimination (RFE) method was used to identify the best predictors (optimal subset of predictors) in the dataset in relation to predicting yield. Subsequently, only these features were retained in the training (Train Optimal) and the testing dataset (Test Optimal). The Train optimal datasets were used to train three separate regression models, one using the random forest (RF) algorithm, one using the gradient boosting machine (GBM) algorithm and one using the extreme gradient boosting (XGB) algorithm. The predictive capabilities of these three models were assessed using the normalized root mean square error (RMSE) metric computed on the associated Test optimal datatsets. Finally, the best regression model, amongst the three, was chosen to compute accumulated local effects (ALE) for predictions made on both the Train Optimal and the Test Optimal datasets and generate yield forecasts for three future time periods under four socioeconomic pathways which were SSP 1-2.6, SSP 2-4.5, SSP 3-7.5 and SSP 5-8.5.

### 2.3 Machine learning algorithms used for feature selection and subsequent forecasting

Three tree-based ensemble ML regression algorithms were used in various ways throughout this work: Random Forest (RF) (Breiman, 2001), Gradient Boosting Machine (GBM) (Friedman, 2001), and XgBoost (XGB) (Chen et al., 2016). Mean decrease of root mean square error (RMSE) calculated by RF was utilized to assess the variable importance of the various predictors in relation their contribution to the overall yield within a Recursive Feature Elimination (RFE) framework (Guyon et al., 2002), and thus mean decrease of root mean square error was useful to select the best predictors in each dataset analyzed. Subsequently, RF, GBM, and XGB were used to train and validate predictive models using only the most important variables identified by RFE in relation to yield (Figure 1). To ascertain overall model performance on the test dataset, Pearson’s correlations between predicted and actual yield values were calculated.

### 2.4 Identifying the influential factors in relation to yield at national and regional scales

Feature selection was used to identify the most influential predictors of overall county-level yield across multiple states (the national model) and within each state (the state-level models). Recursive Feature Elimination (RFE), a backward elimination process implemented in this study by using the R package *caret* (Kuhn, 2008), was used to determine the best predictor subset using the RF algorithm (Breiman, 2001). Subsequent training and validation of predictive models were performed using training and testing datasets containing only the optimal subset of best predictors as deemed by RFE (Figure 1).

### 2.5 Training predictive models and describing relationships between influential predictors and yield at the national and state level

The national training and testing datasets were used to train and validate three separate regression models created with RF, GBM, and XGB (Figure 1). RMSE computed on the testing dataset was used to compare the three models, with the best model selected as the “national model” referred to hereafter. This procedure was repeated for each state to determine the best “state-level model”. To determine which of these models to use for forecasting state-level future yields, the national model was compared to the best state-level model using RMSE. Inputs for national and state-level forecasting models included future projected monthly climate variables as well as the latest county-specific values for areas planted and harvested.

To understand how climate variables drive changes in yield among years, Accumulated Local Effects (ALE) were calculated from model predictions on both the national and state-level training and testing datasets and then plotted (Apley et al., 2020). ALE plots are a robust approach to describing how a predictor variable (climate) impacts a response variable (yield) in the face of dynamic nonlinear relationships where variables are not independent (Molnar, 2021).

An interactive dashboard was developed to facilitate public assessment of modeling outputs [**REDACTED FOR BLINDED REVIEW**]. Cloropleth maps visualize future yield forecasts under the four SSP scenarios, while ALE plots visualize yield responses to each important climate predictor within the national and state-level models. This dashboard was developed using the R package *shiny* (Chang et al., 2022), with visualizations created using the R packages *ggplot2* (Wickham, 2016) and *plotly* (Sievert, 2020).

## 3. RESULTS

There was considerable overlap of predictors deemed important by feature selection at both the national and state level, though with variation in the optimal number of relevant predictors. A total of 37 (out of 38) predictors were deemed to be the most important for accurate yield estimation at the national level (Figure 2), with the most important predictors including maximum temperature and total precipitation in July, as well as minimum temperatures in May and September. Across state-level models (Figures S1-S7), as many as 38 and as few as 4 predictors were retained in the optimal subset, with the most commonly recurring important predictors relating to precipitation and maximum temperatures during the summer peak of the growing season, as well as minimum and maximum temperatures in the spring around planting and the end of the growing season near harvest.

**Figure 2.**
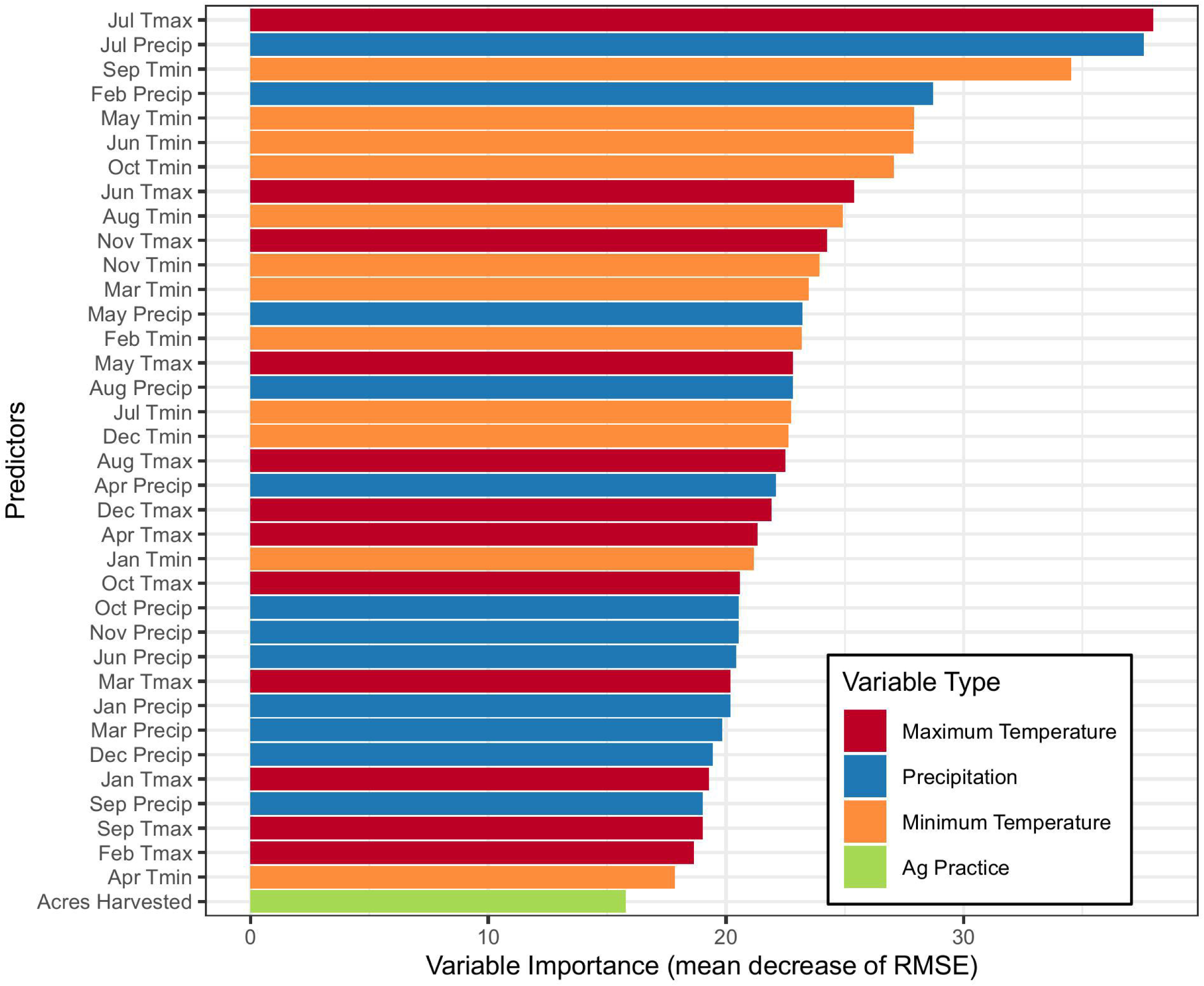
The most important predictors that influence yield at the national level on a relative scale. The metric used for computing variable importance was the mean decrease of root mean square error (RMSE), and the framework used to identify the most important features was the recursive feature elimination (RFE). A random forest algorithm within the framework of RFE was used to create the models and compute variable importance for each predictor variable. The predictors are arranged in order of decreasing importance, where a high value on this scale of relative importance indicates a higher importance for predicting yield.

The best performing national model used the RF regressor (normalized RMSE value of 0.21 and absolute RMSE of 533 lbs/acre) and was thus used to derive future forecasts as well as to calculate ALE values to inspect predictor contributions to yield at the national level. ALE plots suggest that hotter, drier summers result in lower yields, as do overly cool spring and autumn temperatures (Figure 3). July precipitation above 85 mm is sufficient for high yield, while rainfall below this threshold results in a steep reduction of over 100 lbs/acre. Likewise, July maximum temperatures up to 27.5°C are associated with the highest yields, and temperature increases beyond this threshold result in a steep decline in yields of around 200 lbs/acre by 35°C. Minimum temperatures above 8°C in May and 12°C in September are associated with the highest yields, while temperatures below these thresholds are associated with steep declines in yields.

**Figure 3.**
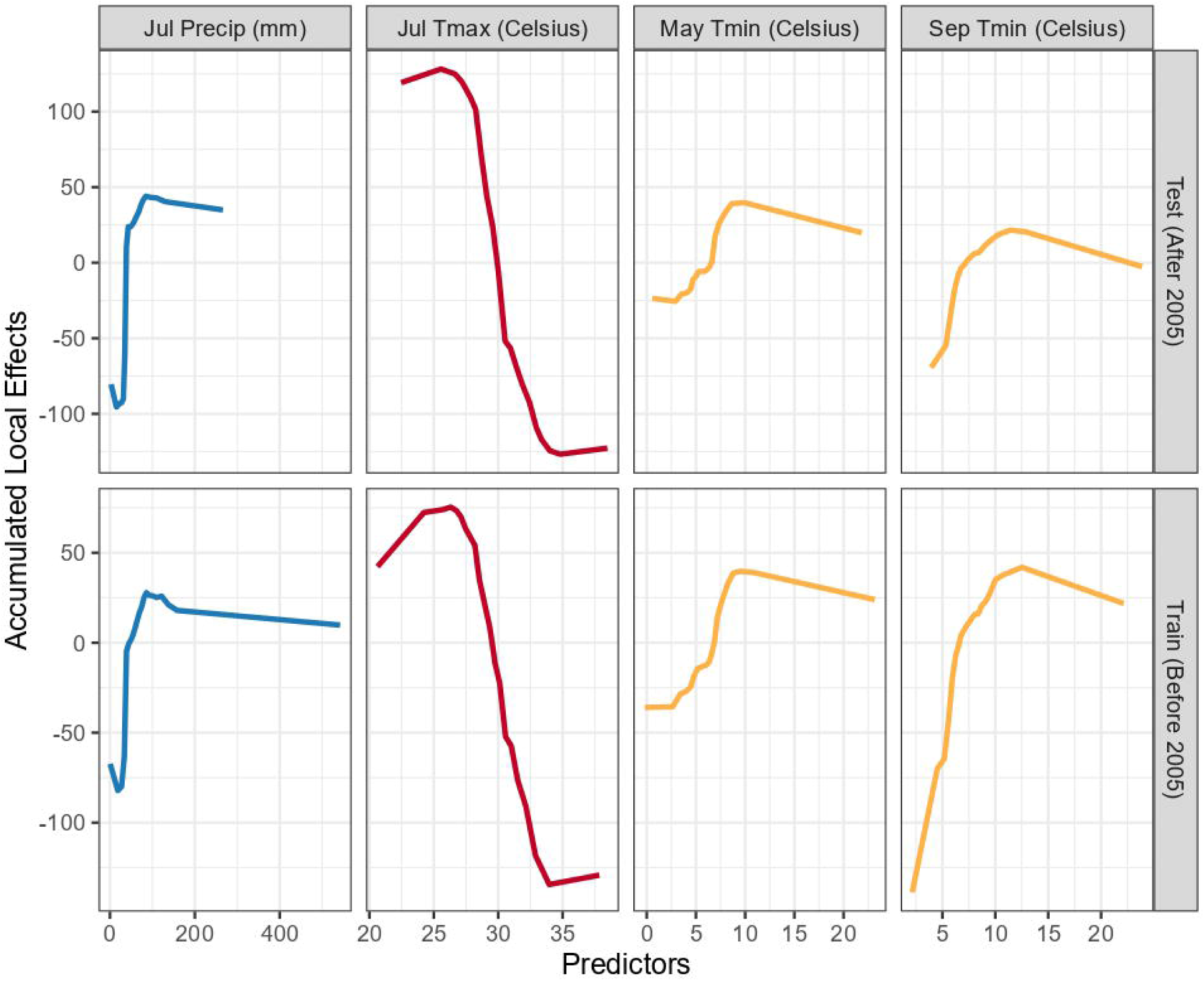
Accumulated Local Effects (ALE) calculated on both training and test datasets for four predictors identified to be important at the national level: total precipitation in July (mm), maximum July temperature (°C), minimum May temperature (°C), and minimum September temperature (°C). The plots articulate the impact of predictor values on yield (in lbs/acre) within the time period modeled.

As the center of sunflower production in the United States, models for North and South Dakota performed similarly to one another and to the national model. The best state level model for South Dakota was the XGB regressor model (normalized RMSE value of 0.25 and absolute RMSE of 542.18 lbs/acre). ALE plots indicate that July maximum temperatures beyond 27°C are associated with a steep decline in yields, as are October minimum temperatures below around 5°C (Figure S8). While an increase in acres planted had a very gradual impact on per-acre yields, the relationship with acres harvested was substantially steeper. This reflects that irrespective of the acreage planted a larger acreage is harvested in South Dakota in favorable years with high per-acre yields, and conversely that fewer acres are taken to harvest in unfavorable years with low per-acre yields. The best state model for North Dakota used the RF regressor (normalized RMSE values of 0.21 and absolute RMSE of 533.59 lbs/acre). As in South Dakota, ALE plots for North Dakota indicate that July maximum temperatures above 27°C are associated with a steep decline in yields (Figure S9), though the training data before 2005 suggest yields also decline when July maximum temperatures are cooler (conditions which do not exist after 2005). Further, July minimum temperatures below 14°C and September minimum temperatures below 7°C are both associated with steep declines in yield. July precipitation below 100mm is associated with severe yield reductions, while increasingly high precipitation (>150mm) is associated with gradual yield declines. Moving southward, state-level models for the central Great Plains began to diverge a bit more from the national model. The best performing state level model for Nebraska was the GBM regressor model (normalized RMSE value of 0.26 and absolute RMSE of 510.7 lbs/acre). Similarly to the Dakotas, ALE plots indicate that early and late season temperatures strongly influence yield in Nebraska (Figure S10). May maximum temperatures below 23°C result in steep yield declines, as do October maximum temperatures below 19°C. In summer, August maximum temperatures above 29°C result in steep yield declines, while August precipitation has a more modest effect with yield declines below approximately 40 mm. The best performing state level model for Kansas was the RF model (normalized RMSE value of 0.22 and absolute RMSE of 389.9 lbs/acre). Much like Nebraska, ALE plots for Kansas indicate that August maximum temperatures above 31°C are associated with steep yield declines, while August precipitation below around 75mm results in modest yield reductions (Figure S11). Further, in Kansas minimum temperatures in February below −4°C result in sharply increasing yields in both the training and testing datasets, with the highest yields in years with the coldest February temperatures – this may indirectly relate to snowfall, pest or pathogen die-off, subsequent planting date, or other unmeasured factors.

Colorado and Minnesota form the western and eastern periphery of major sunflower production, respectively. For both, the best state-level models used the RF regressor (for Colorado normalized RMSE value of 0.19 and absolute RMSE of 392.57 lbs/acre, for Minnesota (0.26 and 383.79 lbs/acre). In Colorado, ALE plots indicate that July precipitation below around 100mm is associated with steep declines in yield (Figure S12). The coolest summers are associated with the highest yields, and increasing August maximum temperatures are associated with sharply declining yields. At the end of the growing season, October minimum temperatures below around 3°C and maximum temperatures below 19°C are associated with lower yields, likely indicative of early frost. In Minnesota, ALE plots indicate that warmer May minimum temperatures above around 6°C increase yield sharply, as do May maximum temperatures above around 18°C up to a peak at around 22°C (Figure S13). Similarly, higher October maximum temperatures sharply increase yields between 11°C and 17°C, but higher end-of-season October precipitation is associated with lower yields. Northern Texas forms the southern limit of commercial sunflower production, and the best state model for Texas used the GBM regressor (normalized RMSE value 0.22 and absolute RMSE value 493.90 lbs/acre). ALE plots indicate erratic responses, with poor agreement between training and test datasets (Figure S14). This may reflect model uncertainty in a state with a lower volume of data available, or shifts in the importance of climatic yield drivers over time.

Assessment of the national model by Pearson’s correlation indicated r=0.32 between predicted yields and actual USDA crop yield statistics on the testing dataset (2005 onwards; Figure 4). The national model on average slightly underpredicted actual yields, with predictions most closely matching actual yields at the high and low ends of predicted yield levels. In North Dakota and South Dakota, the national model outperformed state-level models, so it was used to generate predictions on the state-level test datasets for these two states. The national model predictions correlated well with actual yields in South Dakota (r=0.56), though were less well correlated in North Dakota (r=0.21) and on average underpredicted actual yields in both states. For the remaining states, the respective state-level model outperformed the national model and was used to predict yields for each state testing dataset. The Nebraska model predictions correlated well with actual yields (r=0.48), while models for Texas (r=0.31), Kansas (r=0.31), Colorado (r=0.23), and Minnesota (r=0.16) were less well correlated, though on average did not underpredict actual yields. Low correlations for these four states are likely in part due to the relatively limited yield data available.

**Figure 4.**
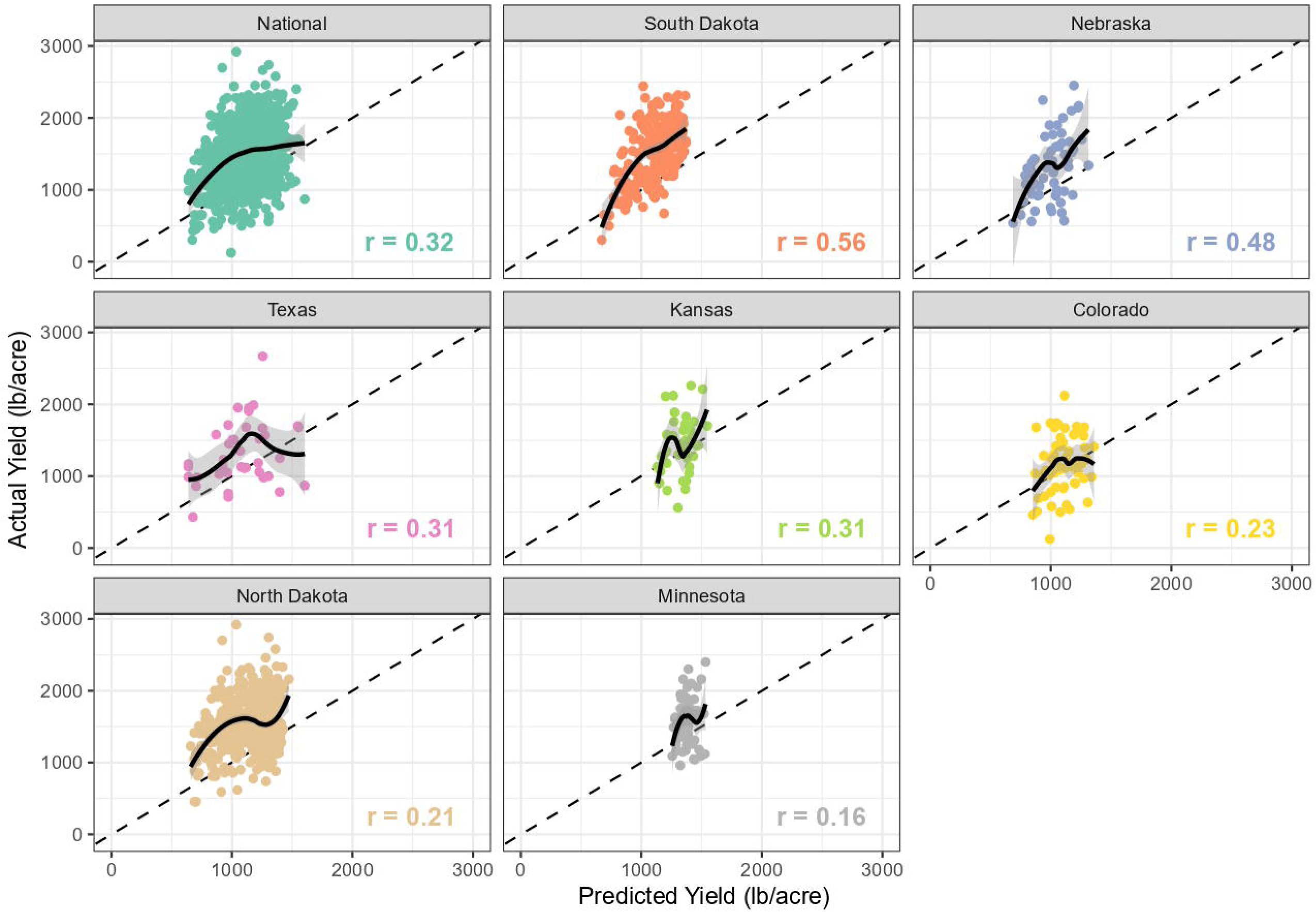
Predictive performance of the best models for a given region (national and state) on the testing dataset using Pearson’s correlation (r) calculated between predicted and actual values. A locally estimated scatterplot smoothing line with a 95% confidence interval describes the relationship between predicted and actual values. In our analysis, we found that the best national model (using the Random Forest regressor) outperformed the state-level models for North Dakota and South Dakota, however, state-level models outperformed the national model for each of the other states. The best state-level models for Colorado, Kansas and Minnesota also used the Random Forest regressor, while for Nebraska and Texas the best state-level models used the Gradient Boosting Machine regressor.

Future yield forecasts at the national level produced similar trends across all four SSPs (Figure 5), with yields sharply declining by about 16% into the first period (2021-2040) relative to the average yield observed between 2000 and 2020, and further modest declines of 1.1-3.5% depending on SSP between the first period and the second period (2041-2060). Average national yields are predicted to remain steady thereafter into the third period (2061-2080). However, a high standard deviation around these estimates of national average yields reflects substantial variation among counties and states in climate change impacts. Among states, the main production regions of North Dakota and South Dakota are predicted to experience 20% and 21% yield declines, respectively, during the first period (2021-2040), with further declines of 1-5% in subsequent decades (Figure S15). Minnesota to the east is similarly predicted to experience 3-4% yield declines during the first period, with negligible changes thereafter. In the central plains Nebraska and Kansas are predicted to experience 0.5-1.5% and 11-12% yield declines, respectively, and single digit percentage increases or decreases thereafter. To the south, Texas exhibits a different pattern, where yields are predicted to decline 21-26% during the first period, and up to 17% more during the second period depending on SSP. Colorado is the only state where yield increases are predicted during the first as well as between the first and the second period, 6-8% during the first period and 1-4% between the first and second period, across all SSPs.

**Figure 5.**
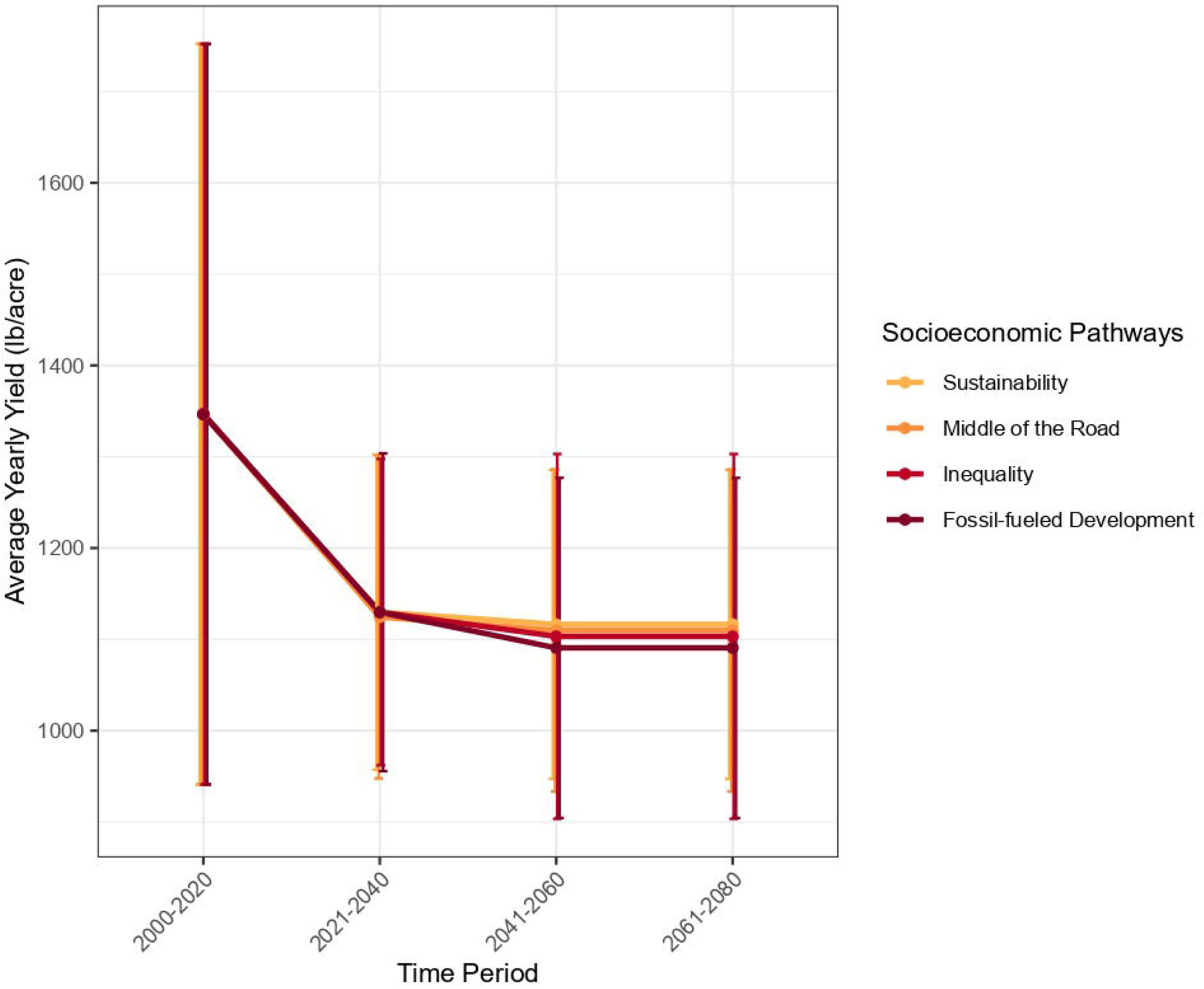
Average yield forecasts at the national level for three time periods and under different Socio-Economic Pathways (SSP). The average annual yield for the year 2022 nationally was approximately 1940 lbs/acre, represented by point estimates and the standard deviation around this mean are represented by the error bars. Future average yield forecasts are also represented as point estimates with the error bars showing variation around these predictions (standard error) and the lines represent change of yield across four time periods and under three socioeconomic pathways (SSP). SSP 1-26, “sustainable future” represents a scenario which involves transitioning towards a more inclusive and environmentally conscious path, focusing on improved global commons management, reduced inequality, and prioritizing human well-being over economic growth, with a shift towards low material consumption and resource efficiency, characterized by an estimated 2.25 °C of Global temperature by 2060 since pre-Industrial age and estimated 38.8 gigatonnes of carbon dioxide emissions. SSP 2-45, “middle of the road”, represents progress towards sustainable development is mixed, with varying levels of economic and social advancements, slow institutional progress, moderate population growth, some environmental improvements, persistent income inequality, and ongoing challenges in reducing vulnerability to societal and environmental changes, characterized by an estimated 2.42 °C rise, and estimated 57.1 gigatonnes of carbon dioxide emissions. SSP 3-75, “regional rivalry”, represents a scenario of increased nationalism, security concerns, and regional conflicts hinder global cooperation, leading to a focus on domestic and regional issues, declining investments in education and technology, slow economic development, material-intensive consumption, persisting or worsening inequalities, varying population growth rates, and severe environmental degradation in some regions due to low international priority on addressing environmental concerns, characterized by an estimated 2.47 °C rise and estimated 68.3 gigatonnes. SSP 5-85, represents a fossil-fueled development scenario, rapid economic growth and technological progress are prioritized through competitive markets, innovation, and investments in human capital, coupled with the exploitation of fossil fuel resources and resource-intensive lifestyles, leading to global economic growth, declining population, successful management of local environmental problems, and reliance on the belief of effectively managing social and ecological systems, including through geo-engineering if needed, and estimated 3.02 °C rise and estimated 101.3 gigatonnes. The climate projections used as inputs for deriving the future yield predictions are from the CMIP6 climate model, the Australian Community Climate and Earth Systems Simulator (ACCESS-ESM1.5) (Ziehn et al., 2020).

## 4. DISCUSSION

Despite rapid growth in the use of ML for agronomic applications, few studies to date incorporate interpretable models to effectively articulate model predictions in an accessible manner (Wolanin et al., 2020; Mateo-Sanchis et al., 2021; Ryo, 2022; Sihi et al., 2022). Demystifying the results from so-called “black box” models can facilitate better understanding of crop growth and development under climatic shifts and under different cultivation practices. As applied here, this approach allowed us to identify relevant relationships between inter-year variation in climate predictors and resulting sunflower yield across seven states. We found that inter-year variation in weather conditions contributes substantially to variation in overall yield at both national and state levels, especially during the typical May-October growing season. Across states, the largest contributors to yield variation are inter-year differences in summer high temperatures (heat waves limiting yield), summer precipitation (drought limiting yield), and overly cool temperatures in the beginning and end of the growing season during planting and establishment (May-June) and maturation and harvest (September-October; Kandel et al., 2020). Interestingly, temperature and precipitation outside of the typical growing season were also important to overall yield dynamics at the national level and in several state models, for example winter precipitation and temperatures (December-February). This indicates that weather conditions outside the growing season can contribute to variability in yield, most likely indirectly through effects on early season soil moisture, timing of soil suitability for planting, and pest and pathogen overwintering success (Unger, 1980; Nielsen, 1998; Gulya et al., 2019; Pantzke et al., 2023). Within the growing season, it has been documented that increases in minimum temperatures cause shifts in phenology across multiple crop species, which in turn influences flowering time and maturity date (Parmesan et al., 2003; Visser et al., 2005; Gordo et al., 2010; Yang et al., 2014). A sustained soil temperature of 10°C during planting is recommended for sunflower germination and establishment (Kandel et al., 2020), and our models reflect that warmer May temperatures (minimums and maximums) are associated with higher yields. In our national model, yield thresholds for minimum May and September temperatures are just above the typical 6°C or 6.7°C growing degree day (GDD) base temperatures for sunflowers (Kiniry et al., 1992; Ferfuia et al., 2015; Kandel et al., 2020). Similarly, multiple experimental studies highlight the optimal temperature range for sunflower growth and development as between 26°C and 29°C (Rondanini et al., 2006; Awais et al., 2017), a pattern which both our national and state-level models detect in maximum temperatures for growing season months with yields declining when temperatures are above this range. As a primarily rain-fed crop, oilseed sunflower requires sufficient precipitation to grow and develop, with suboptimal levels impacting overall yield (Milošević et al., 2015; Kandel et al., 2020). Our national and state-level models all detect precipitation levels in either June, July, or August as one of the most important climatic predictors of yield, typically alongside maximum temperatures for these months, with low precipitation resulting in decreased yields. Water stress is one of the major limiting factors for rainfed sunflower yield, and water limitation combined with heat stress during early flowering up through seed filling negatively impacts yield (Göksoy et al., 2004; Moriondo et al., 2010; García-López et al., 2014). While drought can also be harmful during the vegetative stages, impacting leaf expansion, stem elongation, and mineral nutrient uptake under inadequate soil moisture (Aboudrare et al., 2006; Gunes et al., 2008; Fatemi, 2014; García-López et al., 2014; Hussain et al., 2018), the reproductive stages are especially vulnerable to water limitation due to high transpiration requirements during flowering and seed filling (Lyakh et al., 2014; Totsky et al., 2015). Our national and state-level models not only identify precipitation as important, they describe inflection points below which yield declines sharply (e.g., ∼90mm in July for North Dakota and Colorado, but ∼40mm for South Dakota). Taken together, this example emphasizes how interpretable ML models can capture important environmental thresholds for crop physiological development and quantify impacts on yield accounting for nonlinear interactions with other predictors.

Researchers have predicted future yield decline and stagnation among almost all major crops under a wide range of climate change mitigation strategies (Parry et al., 2004; Ahmed et al., 2016; Iizumi et al., 2018; Yu et al., 2018). Our modeling results predict a comparable decline (16%) in future American sunflower yield over the next two decades under all socioeconomic scenarios explored, with more modest declines thereafter. In the United States, the overall declines in yield expected for major crops like soybean and maize are attributed to increasing temperatures (Basche et al., 2016), along with major changes in cultivation areas for these crops (Wu et al., 2007). Our results for sunflower also suggest that yield declines are likely to be driven by rising temperatures, along with shifts in precipitation.

Globally, substantial variability exists in how future climate and socioeconomic changes will impact different crops across different regions. For example, due to rising temperatures, global maize and soybean yields are projected to plateau even with agronomic adjustments in high-income countries at mid- and high-latitudes, while low-income countries in low latitudes can benefit from mitigation measures to prevent yield stagnation (Iizumi et al., 2018). We also observe regional variability within our models for sunflower, with the largest yield declines in the more productive states of North Dakota, South Dakota, and Kansas, as well as in low productivity states such as Texas. As might be expected, future national forecasts under the most aggressive emissions mitigation (SSP1-2.6) result in smaller yield declines than future forecasts under unmitigated use of fossil fuels (SSP5-8.5), with most states following this general trend. However, differences among SSPs were not especially large, typically only a few percentage points (except for Texas). This indicates that emissions to date are already resulting in climate shifts that will reduce yields during the next two decades, and that mitigation approaches will have a negligible impact on sunflower yields under the existing geographic extent of sunflower cultivation. One exception to the national trend was Colorado, which exhibited yield increases relative to the baseline yield under all four SSPs, indicating that increasing temperatures or shifting precipitation patterns may uniquely benefit Colorado production relative to northern plains states. As noted for other crops, changes in the geographic extent of cultivation may be one strategy to maintain high yields, alongside changing cultivation periods and management practices (Cho and McCarl, 2017; Sloat et al., 2020).

The interpretable ML models generated here consider only one major category of factor that influences crop yields – weather conditions. Like all crops, sunflower productivity is also driven by myriad other factors, including edaphic characteristics (e.g., soil texture, nutrients, water holding capacity, depth to water table), agricultural management practices (e.g., timing of planting and harvesting, planting density, fertilization, irrigation), pest and pathogen outbreaks (and associated interventions), the particular genotype or variety grown (and its inherent yield capacity and stress tolerances), and interactions between all of these factors (de la Vega 2002; Pereira et al., 2019). In particular, it is well documented that different genotypes of sunflower exhibit a wide range of responsiveness to abiotic and biotic stressors (Ouvrard et al., 1996; Panković et al., 1999; Andrianasolo et al., 2016; Hussain et al., 2018). For sunflowers and indeed many crops, comprehensive historical datasets that document these factors simply do not exist, or at nowhere near the temporal or spatial resolution as the county-level historical yield and weather data used here for modeling. Despite this, our models using only climate data as predictors are able to explain over 10% of inter-year variation in yields nationally (r=0.32), and over 23% and 31% of inter-year variation in state-level models for Nebraska (r=0.48) and South Dakota (r=0.56), with agronomically interpretable relationships between predictors and yield. However, these kinds of ML models can certainly be improved by careful incorporation of other types of data that may capture additional yield variation. Soil data may be among the most available, including categorical descriptions of soil types and characteristics through the USDA National Cooperative Soil Survey (e.g, SSURGO; Soil Survey Staff, 2023), or interpolated continuous rasters of estimated soil properties (e.g., SoilGrids; Hengl et al., 2017). Of course, this type of aggregated soil data may not well-reflect conditions within the rooting zone of managed agricultural fields, nor is it well matched to county-level sunflower yield data in the manner that historical weather data is, nor does it capture multi-decade changes in soil conditions. Regardless of these limitations, these kinds of soil data likely provide at least some utility as an estimate of soil impacts on yield across a wide geographic scale. Other data types are less available but could potentially be carefully incorporated with expert domain knowledge. For example, comprehensive disease surveys only date from the early 2000s, with annual or biannual surveys spanning production regions over the past two decades initiated by the National Sunflower Association (Gulya et al., 2019). Coarse estimates of disease prevalence interpolated across the landscape for each year could help capture disease-related inter-year variation in yields, especially if these surveys continue into the future and a larger spatiotemporal dataset can be generated. Organized historical data describing typical county-level agricultural practices (tillage, planting density, fertilization rates, presence of irrigation) or the dominant sunflower variety grown likely do not exist, but general information about regional shifts in the adoption of particular practices or dominant varieties could be incorporated into models as coarse categorical predictors. These kinds of predictors could improve model performance, permit description of the impacts of these practices on yield at a national scale, and increase model utility to extension agents and growers.

To date, there have been few applications of XAI within agronomy, and the broader concept and utility of XAI techniques remain largely unexplored in agriculture (Ryo, 2022). The approach used here in sunflower highlights the utility of XAI for creating interpretable predictive frameworks that can handle nonlinear spatiotemporal relationships common within large agricultural datasets (Bai et al., 2024; Samek et al., 2021). Such frameworks can derive new useful insights from existing historical datasets and can be readily applied to a wide range of crops beyond the handful of most heavily studied commodities. Insights gained for less dominant crops can assist with informing agricultural management decisions for a broader cross-section of growers as the agricultural community navigates the challenges and opportunities of global climate change.

## SUPPLEMENTAL MATERIAL

**Supplemental Methods.** Expanded methodology for statistical modeling.

**Figure S1.** Important yield predictors for South Dakota.

**Figure S2.** Important yield predictors for North Dakota.

**Figure S3.** Important yield predictors for Nebraska.

**Figure S4.** Important yield predictors for Kansas.

**Figure S5.** Important yield predictors for Colorado.

**Figure S6.** Important yield predictors for Minnesota.

**Figure S7.** Important yield predictors for Texas.

**Figure S8.** Accumulated local effects for South Dakota.

**Figure S9.** Accumulated local effects for North Dakota.

**Figure S10.** Accumulated local effects for Nebraska.

**Figure S11.** Accumulated local effects for Kansas.

**Figure S12.** Accumulated local effects for Colorado.

**Figure S13.** Accumulated local effects for Minnesota.

**Figure S14.** Accumulated local effects for Texas.

**Figure S15.** Average yield forecasts for seven states under three climate scenarios.

