## Supplemental Figures for "Sunflower yield modeling with XAI: Historical weather impacts and forecasting"

**Supplementary Methods for [manuscript authors] (2025), *Agronomy Journal***

The following reflects an expanded version of the abridged methods that appear in the main text. This expanded version should be considered the methodology of record.

*2.1. Software and programming language*

Data was compiled, cleaned, and processed using a combination of the R programming language (R Core Team 2023) and ArcGIS Pro software (version 2.9). All data cleaning, visualizations and modeling were performed using various packages created in the R programming environment. All code to replicate the analyses in this study can be accessed through the project GitHub repository. [Link: https://github.com/SamMajumder/Yield_forecast_Sunflowers ]

*2.2. Study area, data sources, and data preparation*

County-level annual "oil type" sunflower seed yield data (measured in pounds per acre) as well as accompanying area planted, and area harvested (both in acres) were acquired programmatically from the United States Department of Agriculture (USDA) National Agricultural Statistics Service (National Agricultural Statistics Service, 2017) using the R package *tidyUSDA* (Lindblad 2022) through an API key. This yield data spanned the temporal range between 1976 and 2022, and included 189 counties across North Dakota, South Dakota, Kansas, Nebraska, Colorado, Minnesota, and Texas. Historical monthly weather data was acquired from the National Oceanic and Atmospheric Administration (NOAA) National Centers for Environmental Information database (NCEI) as multi-layered netCDF raster files containing

5km gridded monthly maximum temperature, minimum temperature, and total precipitation records for the period 1976 to 2022 (Vose et al., 2023).

To obtain future monthly maximum temperature, minimum temperature, and total precipitation projections, multi-layered raster files were obtained from the WorldClim 2.1 repository released in January 2020 (Fick et al., 2017) using the *geodata* R package (Hijmans et al., 2022). WorldClim provides downsampled future climate projections of monthly precipitation, maximum temperatures, and minimum temperatures from a variety of CMIP6 models in various spatial resolutions. We chose to use the 2.5 arc-minute resolution downsampled data (approximately 4.5 km grid cells) to provide similar spatial resolutions for historical and future climate data. The future projections obtained reflected four Shared Socioeconomic Pathways (SSPs; Riahi et al., 2017): SSP 1-2.6, SSP 2-4.5, SSP 3-7.0, and SSP 5-8.5, across three time periods (2021-2040, 2041-2060, and 2061-2080). These projections derive from the CMIP6 climate model, the Australian Community Climate and Earth Systems Simulator (ACCESS-ESM1.5) (Ziehn et al., 2020). SSPs are used to explore future economic and demographic development considering changes in land-use, energy use, and uncertainties regarding greenhouse gas and other air pollutant emissions over the next century (Riahi et al., 2017). SSPs have been implemented in recent reports of the Intergovernmental Panel on Climate Change to ascertain climate goals according to the Paris Agreement announced by the United Nations in 2015 (UNFCCC 2015; IPCC 2022). SSPs are extremely useful when modeling future variation in average global temperature and precipitation patterns due to changing net carbon emissions under various proposed mitigation strategies. The Shared Socioeconomic Pathways (SSPs) outline four scenarios for the future. SSP1 (sustainability) envisions sustainable development with reduced inequality, low resource intensity, and improved global commons management. SSP2 (middle of the road) depicts moderate progress with uneven development, persistent inequality, and slow achievement of sustainable goals. SSP3 (regional rivalry) portrays inequality, nationalism, and environmental degradation. SSP5 (fossil-fueled development) considers rapid economic growth and development using fossil fuels but potential challenges in managing social and ecological systems. Both historical and future weather raster files were first reprojected to the same coordinate system - USA Contiguous Albers Equal Area Conic Projection (Kuniansky 2016; Snyder, 1987) within ArcGIS Pro. This was followed by extracting data from each raster layer at the county centroids (National Weather Service, 1995), followed by storing the extracted data in a dataframe format. Further data preparation was performed within the R programming environment using the packages within the *tidyverse* suite (Wickham et al., 2019), including creating a combined dataset containing the historical weather data alongside historical yield, area planted, and area harvested as well as creating separate dataframes for future climate projections. The most recent area planted, and area harvested data from each county was added to the dataframes containing future climate projections, in order to model impacts on yield with the geographic extent of land under sunflower cultivation held constant. Rows containing missing temperature or precipitation data for specific county-by-year combinations were removed from the combined dataset containing historical yield and weather data, and the dataset was divided into a training and a testing dataset using a temporal cut-off point (Figure 1).

After removing the missing data, the combined dataset now contained a total of 177 counties. Data up to the year 2004 was used for the training dataset, and data from 2005 onwards was used as the testing dataset. Additionally, individual state-level training and testing datasets were also created using the subset of counties from the combined dataset within each of the seven focal states. This resulted in fourteen total state-level datasets, two each (training and testing) from each state. The same temporal cut-off point was used when creating these state-level datasets, and in each case, state-level training datasets contained 70-80% of the total data, with the rest contained in the testing dataset. It was ensured that the counties present in all of the training datasets were also present in the corresponding testing datasets.

*2.3 Machine learning algorithms used for feature selection and subsequent forecasting:*

Three tree-based ensemble ML regression algorithms in various ways throughout this work: Random Forest (RF) (Breiman, 2001), Gradient Boosting Machine (GBM) (Friedman, 2001), and XgBoost (XGB) (Chen et al., 2016). Mean decrease of root mean square error calculated by RF was utilized to calculate the variable importance of the various predictors in relation their contribution to the overall yield within a Recursive Feature Elimination (RFE) framework (Guyon et al., 2002), and thus mean decrease of root mean square error was useful to select the best predictors in each dataset analyzed. Subsequently, RF, GBM, and XGB were used to train and validate predictive models using only the most important variables identified by RFE in relation to yield. These predictive models are suitable for both examining historical climate contributions to yield variation as well as creating forecasts over future timeframes, bootstrapped version of the dataset, created by random sampling without replacement, is used to train decision trees within the RF ensemble (Pal 2005; Valletta et al., 2017). The predictive capability of the trained ensemble is tested using the portion of the data not included in this bootstrapped version, called the “out of bag” (OOB) data (Cutler et al., 2012; Valletta et al., 2017). GBM, a “boosting algorithm”, reduces the error with each subsequent decision tree in the ensemble, and in this way improves the prediction accuracy of the overall model in a sequential manner (Friedman, 2001).

XGB, a more regularized and optimized version of GBM, uses the same principles, except L2 (Ridge regularization) and L1 regularizations (Lasso regularization) are applied to prevent overfitting. Both regularizations use penalty terms to achieve this, however, L1 regularization adds the absolute value of the coefficient as a penalty term and L2 adds the squared value of the coefficient. Furthermore, target oriented-cross validation (TOCV) (Meyer et al., 2018) was used whenever a tree-based algorithm was applied to the training dataset in order to maintain the spatiotemporal structure of the data and minimize the chances of overfitting. TOCV was implemented in the analyses using the CAST package within the R environment (Meyer et al., 2022). The metric used to quantify and evaluate model performance was root mean square error (RMSE) which is the average distance between the predicted value from a regression model and the actual value in the dataset. Additionally, the absolute RMSE values from each model were normalized between 0 and 1 to facilitate interpretability. This was achieved by dividing the absolute RMSE values with the range of the response (yield) value. To ascertain overall model performance on the test dataset, Pearson's correlations between predicted and actual yield values were calculated.

### *2.4 Identifying the influential factors in relation to yield at national and regional scales.*

Feature selection was used to identify the most influential predictors (optimal number of relevant predictors) in relation to the overall county-level yield across multiple states (the national model) and within each state (the state-level models). Feature selection is not only necessary for better model performance in regard to prediction, but it also facilitates better understanding of the fundamental mechanisms within a biological system that influence the outcome of interest (Guyon et al., 2002). In ML literature, "features" refers to predictors and in this study, performing a feature selection step was necessary to identify the core subset of climate variables that influence annual yield. Recursive Feature Elimination (RFE), a backward elimination process implemented in this study by using the R package *caret* (Kuhn, 2008), was used to determine the best predictor subset. In the context of this study, this subset contained the specific set of predictors which are the most influential to yield prediction. Within the RFE framework, multiple models are trained on the dataset using a regression algorithm chosen by the user (Guyon et al., 2008) and in our case we used RF (Breiman, 2001). At each step after training using TOCV (Meyer, et al. 2018), importance values were calculated for each predictor to ascertain its contribution to the prediction of yield. The predictors were ranked based on their importance and the predictor with the least importance value was eliminated, with this process repeating until no variables remained in the model. This method of feature selection was applied to the national model training dataset (containing data from all seven states) to uncover the core subset of important variables relevant to yield prediction nationwide, as well as to state-level training datasets to potentially detect predictors relevant only within a given state. The metric used to calculate and rank variable importance was Mean Decrease of Root Mean Square Error (Breiman, 2001). Subsequent training and validation of predictive models were performed using training and testing datasets containing only the subset of best predictors as deemed by RFE (Figure 1).

### *2.5 Training predictive models and describing relationships between influential predictors and yield at the national and state-level.*

The national training and testing datasets (containing data from all seven states) were used to train and validate three separate regression models: one created with RF, one with GBM and one with XGB. The performance of these models was compared using RMSE computed on the testing dataset, and the best model among the three was determined to be the “national model” referred to hereafter. This procedure was repeated for each state, with the three regression models trained (RF, GBM and XGB), validated, and compared to determine the best “state-level model” for that particular state. To determine whether to use a state-level model or the national model when forecasting future yields in each state, the national model was compared to the best performing state-level model, with the model with the lowest RMSE value (both absolute and normalized) selected to perform forecasting. The future projections for total monthly precipitation, maximum and minimum temperature as well as latest county specific values for areas planted and area harvested, were used as inputs to the best forecasting models, when deriving future county-wide yield forecasts for both the nation and each state.

The national and state-level models were also used to understand how important climate variables (predictors) drive changes in yield within the historical testing and training datasets. To achieve this, Accumulated Local Effects (ALE) were calculated from model predictions on both the national and state-level training and testing datasets and then plotted (Apley et al., 2020). The values of a predictor variable of interest are divided into different intervals and the difference in prediction at each interval is recorded and accumulated, thus creating an ALE curve (Molnar, 2021; Apley et al., 2020). ALE plots describe the manner in which values of a predictor variable (here, climate) impact the value of the response variable (here, yield), and enabled extrapolation of the dynamic nonlinear relationships between predictor variables and the predicted yield. ALE plots are robust when variables in a dataset are correlated with each other, as common for climate variables, as ALE does not assume independence of variables (Molnar, 2021). ALE values for important variables at the national level were derived from model predictions made by the “national model” on both the training and the test dataset. Each state-level ALE values were calculated from predictions made by the best “state-level model” for that given state. Computing ALE on both the training and test datasets enabled us to view and compare patterns between two temporal scales i.e., up to 2004, and after 2005. An interactive dashboard was developed to visualize future yield forecasts under four SSP scenarios using choropleth maps. Additionally, ALE plots visualizing thresholds for each important predictor within the national model and state-level models are also presented in this dashboard (Link: https://sammajumder.shinyapps.io/SunScope/ ). The dashboard was developed using the R package *shiny* (Chang et al., 2022) and the interactive visualizations on this dashboard (as well as static figures presented here) were created using the R packages *ggplot2* (Wickham, 2016) and *plotly* (Sievert, 2020).

IPCC, 2022: Climate Change 2022*:* Impacts, Adaptation, and Vulnerability Contribution of Working Group II to the Sixth Assessment Report of the Intergovernmental Panel on Climate Change. Page 3056 [H.-O. Pörtner, D.C. Roberts, M. Tignor, E.S. Poloczanska, K. Mintenbeck, A. Alegría, M. Craig, S. Langsdorf, S. Löschke, V. Möller, A. Okem, B. Rama (eds.)]. Cambridge University Press. Cambridge University Press, Cambridge, UK and New York, NY, USA.

Kuhn M 2008 Building predictive models in R using the Caret package. Journal of Statistical Software 28: 1–26.

Kuniansky EL 2017 Custom map projections for regional groundwater models. Groundwater 55(2): 255-260.

Lindblad B 2022 tidyUSDA: A Minimal Tool Set for Gathering USDA Quick Stat Data for Analysis and Visualization. R package version 0.4.0.

Meyer H, C Reudenbach, T Hengl, M Katurji, T Nauss 2018 Improving performance of spatio- temporal machine learning models using forward feature selection and target-oriented validation. Environmental Modelling & Software 101: 1-9.

Meyer H., M Ludwig 2022 CAST: 'caret' Applications for Spatial-Temporal Models. R package version 0.6.0.

Molnar C 2022 Interpretable Machine Learning: A Guide for Making Black Box Models Explainable (2nd ed.).

National Weather Service. (1995). *Counties of U.S.* [Vector digital data]. Silver Spring, MD: National Weather Service.

Pal M 2005 Random Forest Classifier for Remote Sensing Classification. International Journal of Remote Sensing 26: 217–22.

R Foundation for Statistical Computing. 2022. Version 4.2.1 (2022-06-23). The R Foundation for Statistical Computing, Vienna Austria. Website https://www.R-project.org/ [accessed March 2023].

Riahi, K, DPV Vuuren, E Kriegler, J Edmonds, BCO Neill, S Fujimori, N Bauer et al., 2017 The Shared Socioeconomic Pathways and their energy, land use, and greenhouse gas emissions implications: An overview. Global Environmental Change 42:153-168.

Sievert C 2020 Interactive Web-Based Data Visualization with R, plotly, and shiny. Chapman and Hall/CRC Florida

Snyder JP 1987 Map projections--A working manual. US Government Printing Office 1395.

Soil Survey Staff Natural Resources Conservation Service United States Department of Agriculture. Soil Survey Geographic (SSURGO) Database. Available online at https://sdmdataaccess.sc.egov.usda.gov. Accessed [6/9/2023].

United Nations Framework Convention on Climate Change (UNFCCC) 2015 The Paris Agreement. UNFCCC, New York.

USDA National Agricultural Statistics Service. (2017). *Census of Agriculture*. U.S. Department of Agriculture.

Valletta JJ, C Torney, M Kings, A Thornton, J Madden 2017 Applications of Machine Learning in Animal Behaviour Studies. Animal Behaviour 124: 203–20.

Vose SR, S Applequist, M Squires, I Durre, M Menne, WJ Matthew, N Claude Jr., C Fenimore, K Gleason,D Arndt 2014 NOAA Monthly U.S. Climate Gridded Dataset (NClimGrid), Version 1. [subset used 1976-2022]. NOAA National Centers for Environmental Information. DOI:10.7289/V5SX6B56 [Accessed on March 2023].

Wickham H 2016 ggplot2: Elegant Graphics for Data Analysis. Springer-Verlag New York. ISBN 978-3-319-24277-4.

Wickham H, M Averick, J Bryan, W Chang, LDA McGowan, R François, G Grolemund et al., 2019 Welcome to the Tidyverse. Journal of Open Source Software 4: 1686.

Ziehn, T., Chamberlain, M. A., Law, R. M., Lenton, A., Bodman, R. W., Dix, M., & Stevens, L. (2020). The Australian earth system model: ACCESS-ESM1.5. *Journal of Southern Hemisphere Earth Systems Science, 70*(1), 193-214.
